# Deciphering cell lineage specification during male sex determination with single-cell RNA sequencing

**DOI:** 10.1101/190264

**Authors:** Isabelle Stévant, Yasmine Neirjinck, Christelle Borel, Jessica Escoffier, Lee B. Smith, Stylianos E. Antonarakis, Emmanouil T. Dermitzakis, Serge Nef

**Affiliations:** Department of Genetic Medicine and Development, University of Geneva, 1211 Geneva, Switzerland; iGE3, Institute of Genetics and Genomics of Geneva, University of Geneva, 1211 Geneva, Switzerland; SIB, Swiss Institute of Bioinformatics, University of Geneva, 1211 Geneva, Switzerland; MRC Centre for Reproductive Health, University of Edinburgh, Edinburgh EH16 4TJ, UK; School of Environmental and Life Sciences, University of Newcastle, Callaghan, NSW 2308, Australia.

**Keywords:** Single-cell RNA-Seq, sex determination, testis, Sertoli cell, fetal Leydig cell, progenitors, differentiation, lineage specification, cell fate decision, gene expression

## Abstract

The gonad is a unique biological system for studying cell fate decisions. However, major questions remain regarding the identity of somatic progenitor cells and the transcriptional events driving cell differentiation. Using time course single cell RNA sequencing on XY mouse gonads during sex determination, we identified a single population of somatic progenitor cells prior sex determination. A subset of these progenitors differentiate into Sertoli cells, a process characterized by a highly dynamic genetic program consisting of sequential waves of gene expression. Another subset of multipotent cells maintains their progenitor state but undergo significant transcriptional changes that restrict their competence towards a steroidogenic fate required for the differentiation of fetal Leydig cells. These results question the dogma of the existence of two distinct somatic cell lineages at the onset of sex determination and propose a new model of lineage specification from a unique progenitor cell population.

## Introduction

Testis development is a powerful model for the study of cell lineage specification and sex-specific cell differentiation. Prior to sex determination, the gonadal ridge is composed of primordial germ cells and uncharacterized somatic precursor cells. These precursors originate primarily from the overlying coelomic epithelium (Svingen and Koopman, 2013) and communally express the transcription factors NR5A1 (nuclear receptor subfamily 5 group A member 1, also known as SF1 and Ad4BP) (Hatano et al., 1996; Luo et al., 1994), WT1 (Wilms' tumour (WT) suppressor 1, GATA4 (GATA transcription factor 4) and LHX9 (Lim homeobox gene 9) (Birk et al., 2000; Mazaud et al., 2002). During gonadal ridge formation, proliferating NR5A1+ cells delaminate from the coelomic epithelium, enter the gonad and commit to the supporting cell lineage or to the steroidogenic cell lineage which will give rise to respectively Sertoli and fetal Leydig cells after male sex determination (DeFalco et al., 2011; Karl and Capel, 1998). Sertoli cells are the first somatic cell type to differentiate around embryonic day 11.5 (E11.5) in the mouse (Albrecht and Eicher, 2001; Koopman et al., 1990) and are essential to coordinate testis development. Sertoli cells aggregate around germ cells, forming structures named testis cords, and orchestrate the differentiation of other somatic cells including fetal Leydig cells by E12.5 (Habert et al., 2001). The increase in fetal Leydig cell number between E12.5 and E15.5 occurs by recruitment and differentiation of interstitial Leydig progenitor cells rather than by mitotic division of differentiated fetal Leydig cells (Barsoum and Yao, 2010a; Migrenne et al., 2001; Miyabayashi et al., 2013; Wen et al., 2016).

Yet, it remains unclear whether Sertoli and fetal Leydig cells derive from a single or from two predefined precursor populations, the supporting and the steroidogenic cell lineages. A better characterization of cell heterogeneity of the bipotential gonads prior to sex determination has been hampered by the absence of specific markers for precursor populations. A more complete understanding of lineage specification during testis development is critical to identify the main regulators of normal testicular development and tissue homeostasis.

Existing time course transcriptomic assays using sample pools of purified testicular cells has provided valuable evidence of the dramatic changes occurring during cell differentiation (Jameson et al., 2012; Munger et al., 2013; Nef et al., 2005), but a higher resolution is needed to evaluate cell type heterogeneity and the precise dynamics of gene expression during cell lineage specification.

In this study we utilized single cell RNA sequencing on NR5A1+ cells to perform an unsupervised reconstruction of somatic cell lineage progression in the developing testis, prior to, during and after sex determination. At E10.5, in the bipotential gonads, we observed a single uncommitted somatic cell population. This multipotent progenitor cell population first give birth to Sertoli cells around E11.5, which is driven by a rapid and highly dynamic differentiation program. Then, the remaining progenitors evolve transcriptionally to express steroidogenic precursor cell markers at the origin of Leydig cells.

## Results

### Somatic cell purification and single-cell RNA-sequencing

To isolate and RNA-sequence individual somatic cells of the gonads prior, during and after sex determination, we collected gonads from *Tg(Nr5a1-GFP)* male mice at five different developmental stages of testicular differentiation (E10.5, E11.5, E12.5, E13.5, E16.5) (Nef et al., 2005; Pitetti et al., 2013; Stallings, 2002). *Nr5a1* is specifically expressed in the somatic cells of the developing gonads as early as E10.5, in Sertoli cells, fetal Leydig cells, and interstitial cells (**Figure 1A-B, Figure S1**).

**Figure 1:**
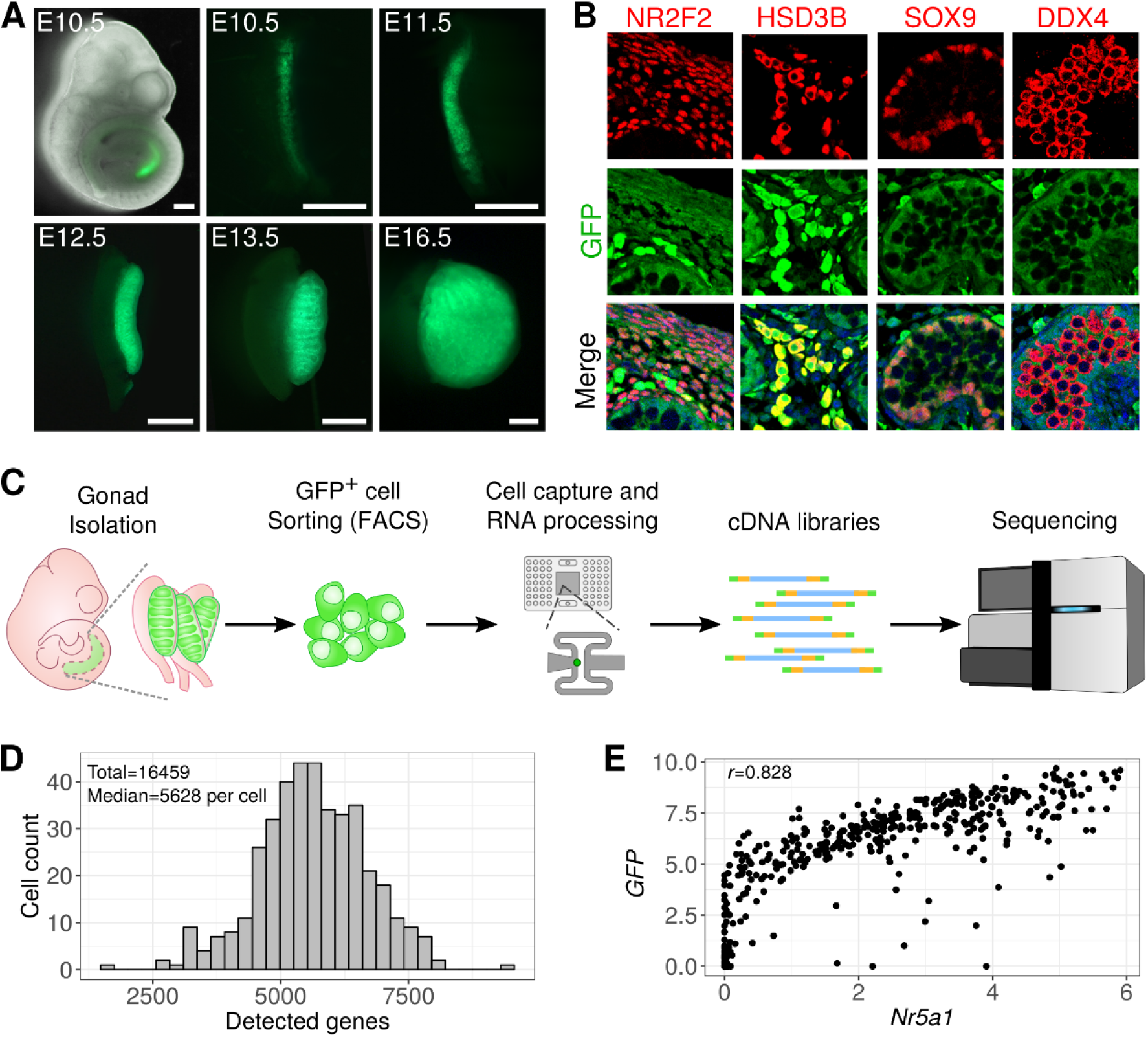
Experimental protocol. (A) Images of E10.5 whole *Tg(Nr5a1-GFP)* mouse embryo (merge of bright field and UV light) and XY gonads at five stages of development under UV light (scale bars=100μm). (B) Co-immunofluorescence of GFP and marker genes for interstitial progenitors (NR2F2), fetal Leydig cells (HSD3B), Sertoli cells (SOX9) and germ cells (DDX4) respectively at E16.5. GFP co-localizes with the somatic cell markers but not with the germ cell marker. (C) Experimental design. XY gonads at each stage were collected, Nr5a1-GFP+ cells were FACS-sorted and captured, harvested cDNA processed for libraries and sequenced. (D) Distribution of the number of detected genes per cell. (E) Correlation between the expression of the *Nr5a1-GFP* transgene and the endogenous *Nr5a1* gene.

The NR5A1-GFP+ cells from the genital ridges were FACS-sorted and individually captured and processed with the Fluidigm C1 Autoprep System (**Figure 1C, Figure S1**). We prepared and sequenced 435 single-cell libraries, from which 400 passed the quality filters (**Table S1, Figure S2 & S3, and Supplemental experimental procedures**). Cells expressed a total of 16 459 protein-coding genes (RPKM>0) with a median of 5628 genes per cell (**Figure 1D**). A strong correlation (r=0.82, Spearman correlation) between the levels of *GFP* and endogenous *Nr5a1* transcripts confirmed the reliability of the Tg(Nr5a1-GFP) reporter (**Figure 1E**).

### Single-cell transcriptomics identifies six gonadal somatic cell populations

To identify the different cell populations captured in our experiments, we selected the highly variable genes and performed a hierarchical clustering on the significant principal components, and we visualized the cell clusters with t-SNE (**Supplemental experimental procedures**). We obtained six cell clusters, each mixing different embryonic stages (**Figure 2A&B**). The dendrogram (**Figure 2C**) reveals the relative distances between the cell clusters. Cluster 1 (C1) is the most distant, and C2 and C3 the most similar. We found 2802 differentially expressed genes (*q-value*<0.05) between the six cell clusters (**Supplementary Data 1**). GO terms of the cluster up-regulated genes (**Figure 2D, Supplementary Data 2**) associate C1 to angiogenesis, C2 and C3 to Wnt signaling and process and urogenital system development, C4 to sex differentiation, C5 to steroid metabolic process, and C6 to cell cycle.

**Figure 2:**
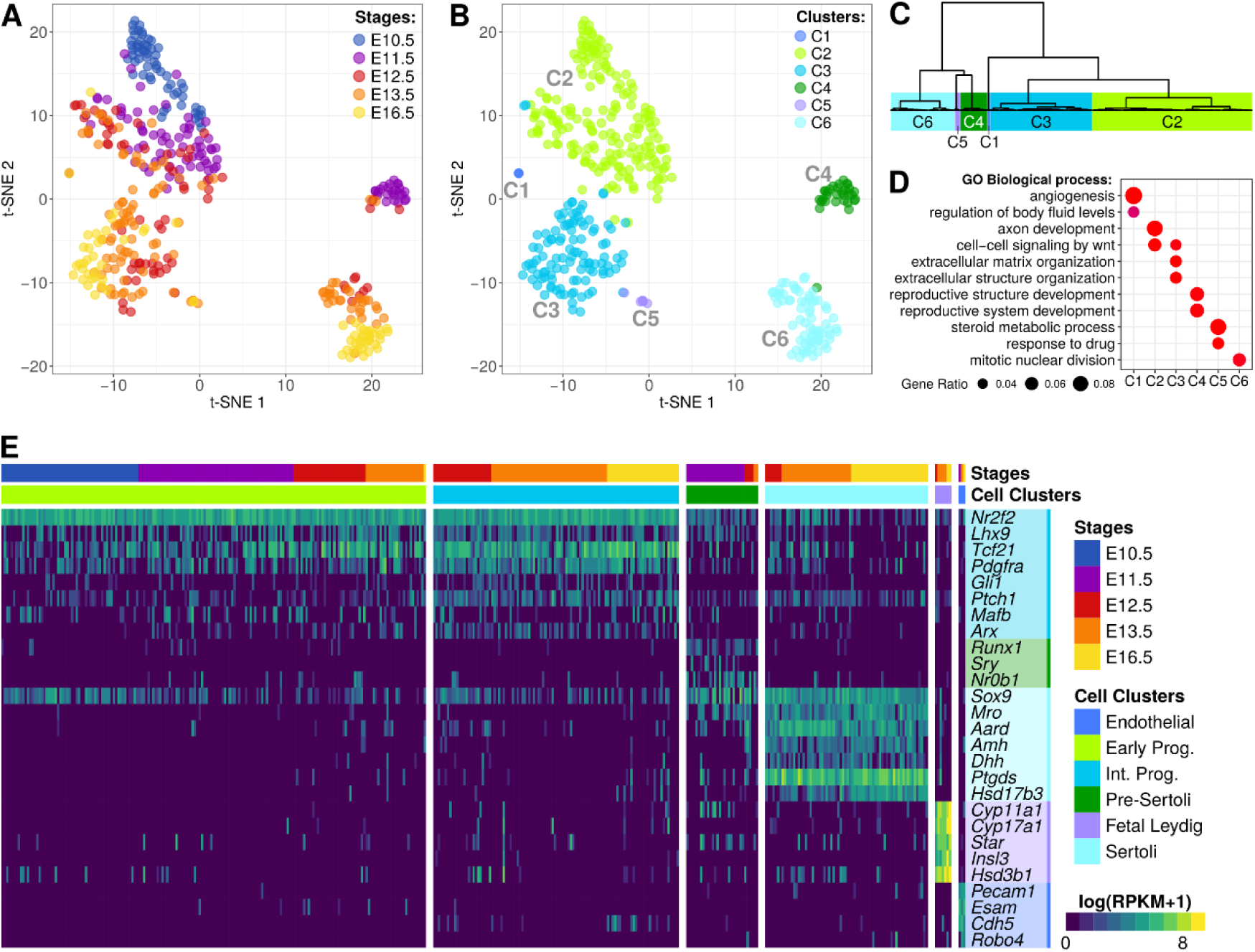
Identification of the cell populations. (A) and (B) Two-dimensional t-SNE representation of 400 GFP+ single-cell transcriptomes on the most variable genes. Cells are colored by embryonic stages (A) and by cell clusters (B). (C) Dendrogram of the hierarchical clustering on the principal components (HCPC). (D) GO biological process associated with the up-regulated genes from each of the six cell clusters. (D) Heatmap of the expression of marker genes of testicular somatic cell types per cell clusters.

The embryonic stages together with significant enrichment of co-expressed marker genes of the different testicular cell populations allow us to confidently assign an identity to each of the cell clusters (**Figure 2E, Figure S4**). C1 is composed of 3 cells from E11.5, E13.5 and E16.5 expressing endothelial cell marker genes, consistently with the associated GO terms. C2 contains 183 cells from E10.5 to E13.5 It is the only cell cluster containing the E10.5 cells and is highly similar to C3. Therefore we associate C2 cells to the early somatic progenitor cells. C3 contains 106 cells from E12.5 to E16.5 and is enriched in interstitial progenitor marker genes while C4 is composed of 31 cells from E11.5 to E13.5 expressing pre-Sertoli cell markers. C5 contains 7 cells from E12.5, E13.5 and E16.5 expressing fetal Leydig cell markers and finally C6 is composed of 70 cells from E12.5 to E16.5 and is enriched in Sertoli cell marker genes.

Overall, with our single-cell RNA-seq experiments on Nr5a1-GFP+ cells, we captured and identified six somatic cell populations in developing testis. We detected a single progenitor cell population from E10.5 that remains until E13.5 in the developing testis, the committed Pre-Sertoli cells at E11.5, the differentiating Sertoli cells from E12.5, the interstitial progenitor cells from E12.5, and the fetal Leydig cells from E12.5. We also detected three endothelial cells although they are not known to be Nr5a1+ cells.

### Pseudotime reconstruction identifies cell lineage specification from the E10.5 cells

To understand the lineage specification from the E10.5 progenitor cells and identify the mechanisms controlling cell fate decision, we performed an *in silico* reconstruction of the cell lineages using diffusion map and ordered them along a pseudotime (**Figure 3A-B, Supplementary experimental procedures**). We obtained three cell lineages, the interstitial progenitor cell lineage, the Sertoli cell lineage and the fetal Leydig cell lineage, in accordance to the previous cell clustering. These three lineages share a common origin starting from the E10.5 cells. No specific cell lineage was found for the endothelial cells which were assigned to the interstitial progenitor cell lineage.

**Figure 3:**
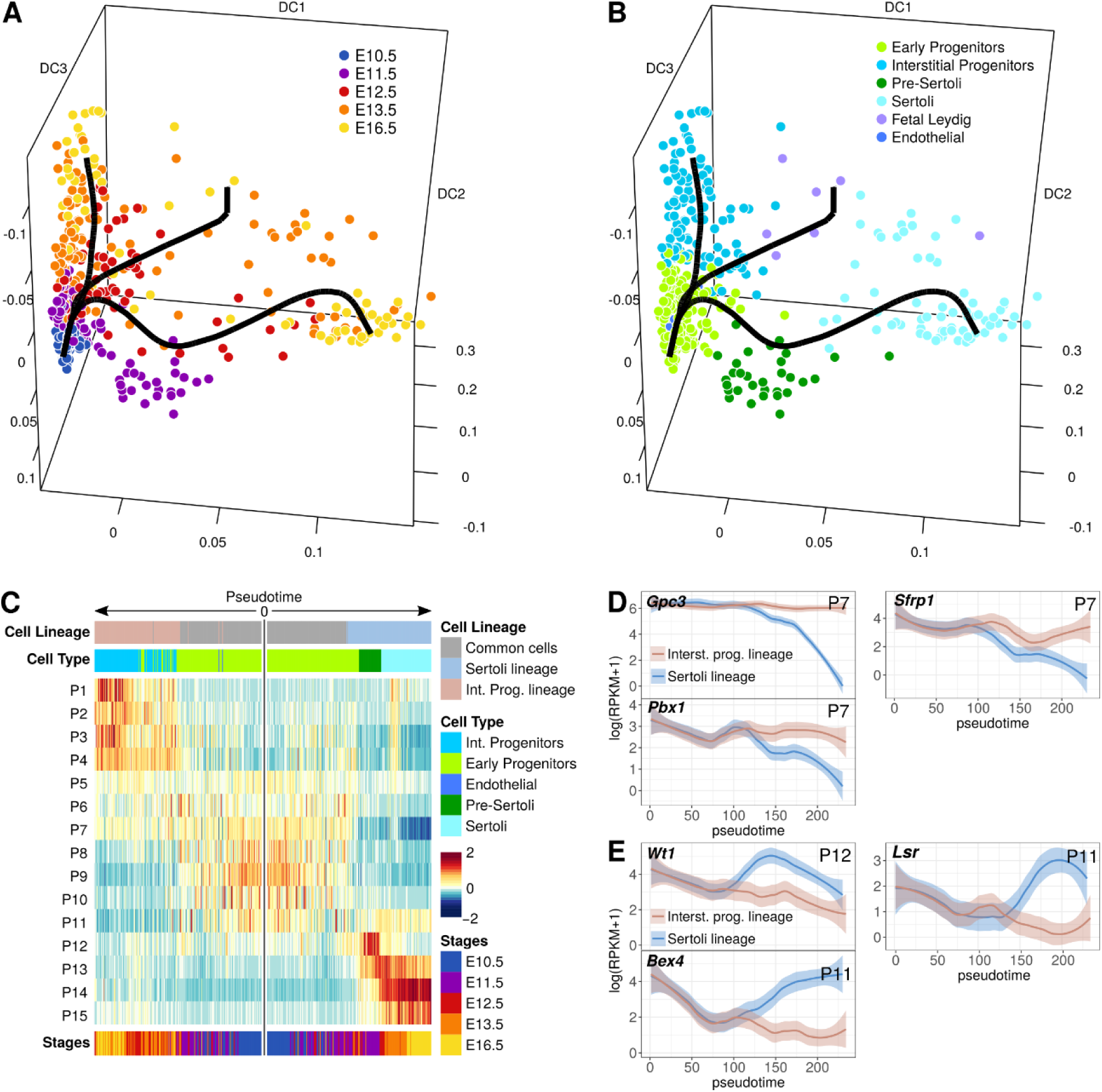
Cell lineage and pseudotime prediction shows different cell differentiation strategies. (A) and (B) Diffusion map on the most variable genes and reconstruction of the cell lineages with Slingshot. Each dot represent a cell and the black lines represent the estimated cell lineages. (A) is colored by embryonic stages and (B) by cell clusters. (C) Heatmap representing the transcriptomic progress of the progenitor and the Sertoli cell lineages specific genes. Genes were grouped by expression profiles with k-means and the heatmap shows the normalized average expression of each group. Cells common to both lineages are located in the middle of the heatmap, and progression of each lineage are presented through both extremities, with progenitors on the left and Sertoli cells on the right side. (D) and (E) Smoothed expression profiles of interstitial progenitor (D) and Sertoli lineage (E) specific genes. The solid line represents the loess regression, and the fade band is the 95% confidence interval of the model.

To compare how the lineages acquire their identity, we selected cell cluster-enriched genes (differentially expressed genes with q-value<1e10^−5^) and looked at their expression dynamics along the predicted pseudotime from their common origin at E10.5 until E16.5 (**Figure 3C**). The small number of fetal Leydig cell prevent a high resolution understanding of their differentiation process (not shown).

The comparison of the interstitial progenitor and the Sertoli cell lineage (**Figure 3C, Supplementary Data 3**) reveals that 45% (162 out of 357 genes, gene profiles P5, P6 and P7) of the interstitial progenitor enriched genes are also expressed in the common progenitor cells. GO terms associated with these genes are relative to urogenital system development and epithelium development (**Supplementary Data 4**). Among these genes, we found regulators of the Wnt signaling pathway, such as *Gpc3* and *Sfrp1* (**Figure 3D**). *Gpc3* modulates IGF2 interaction with its receptor and has been described as responsible for the X-linked recessive Simpson-Golabi-Behmel syndrome in human. Patients display genital abnormalities including testicular dysplasia and cryptorchidism (Griffith et al., 2009). *Sfrp1* is a secreted negative regulators of the Wnt signaling pathway which have been identified as important for testis descent and normal testicular development (Warr et al., 2009). We also found *Pbx1*, which have been previously described as expressed in the interstitial compartment of the testis and is required for a normal urogenital differentiation (Schnabel et al., 2003). The other interstitial progenitor specific genes display a gradual over-expression in time from E12.5 to E16.5 (P1 to P4).

On the other hand, 22% (**Figure 3C**, 75 out of 340 genes, gene profiles P11 and P12) of the Sertoli cells enriched genes are also expressed in the common progenitor cells. Among them we found *Wtl* (**Figure 3E**), which is necessary for genital ridge formation and Sertoli cell differentiation and maintenance, *Bex4* (brain expressed X-linked 4), which is expressed in Sertoli cells but the function is poorly understood (Yu et al., 2017), and *Lsr* (lipolysis stimulated lipoprotein receptor), which the function is unknown in testis development. The other Sertoli cell specific genes display a sharp and strong overexpression at the exact moment of cell differentiation (P13 to P15).

The lineage reconstruction and the predicted pseudotime allow us to accurately describe the genetic program driving cell differentiation. With this analysis, we showed that the lineage specification of the interstitial progenitor cells display linear and smooth processes, by slowly increasing the expression of their specific genes. In contrast, Sertoli cell lineage differentiates from the common progenitor cells by downregulating the progenitor genes and rapidly upregulating their specific genes.

### Progenitor cells gradually acquire a steroidogenic fate

To deeper evaluate the transcriptional changes of these progenitor cells from E10.5 to E16.5, we performed a differential expression analysis as a function of the pseudotime on the whole transcriptome (**Supplementary experimental procedures**). We identified 1734 genes presenting a significant change through the pseudotime (*q-value*<0.05) and classified them into nine expression profiles (P1 to P9, **Figure 4A, and Supplementary Data 5**).

**Figure 4:**
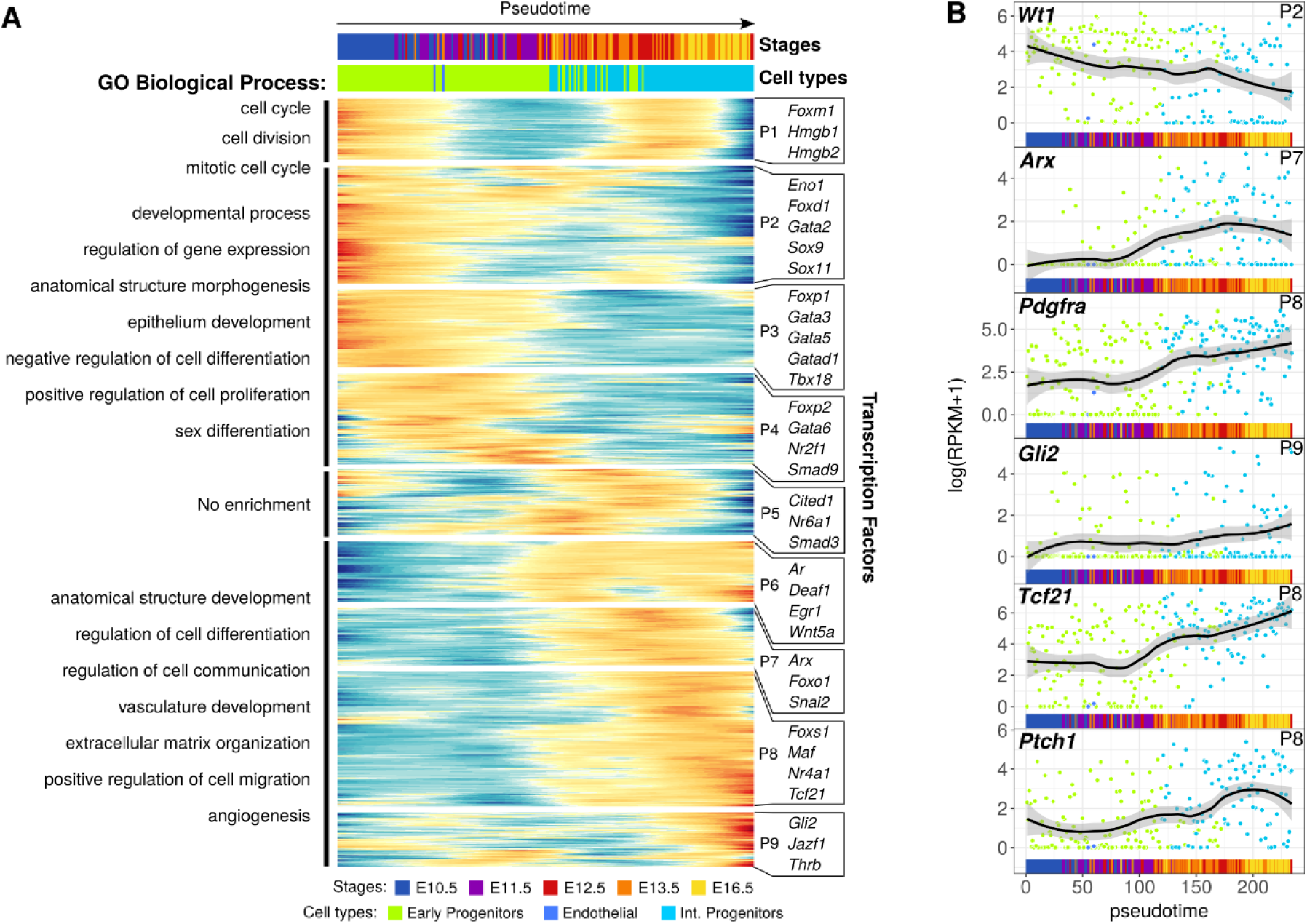
Dynamics of the progenitor transcriptome from E10.5 onward. (A) Heatmap of the normalized smoothed expression of genes differentially expressed as a function of the pseudotime. Genes were grouped by expression profiles with hierarchical clustering. GO terms from enrichment analysis reveals the biological processes of the genes according to their expression profile. On the right are listed some of the transcription factors expressed in each expression profile. (B) Expression profiles of relevant genes reflecting the acquisition of a steroidogenic fate. Each dot represent a cell, the solid line represents the loess regression, and the fade band is the 95% confidence interval of the model.

The expression profile P1 shows genes that are expressed twice during the cell development, once at E10.5 and once around E12.5-E13.5. These genes are related to mitotic cell division.

The expression profiles P2 to P4 present genes expressed between E10.5 and E11.5 and are related to developmental process, regulation of gene expression, and epithelium development, consistently with the cellular identity of the NR5A1^+^ cells of the genital ridge before sex determination. These three expression profiles also contains genes related to stem cell maintenance and renewal related genes *(Sall1, Lin28a, Trim71)* (Basta et al., 2014; Cuevas et al., 2015; Yang et al., 2015), the negative cell differentiation regulator *Foxp1* (van Keimpema et al., 2015; Li et al., 2012; Takayama et al., 2008), and *Wt1* and *Cbx2* necessary for *Sry* expression and thus Sertoli cell differentiation (Katoh-Fukui et al., 2012; Wilhelm and Englert, 2002) (**Supplementary Data 5**).

Expression profiles P6 to P9 contain genes expressed from E12.5 onward. Associated GO terms are related to extracellular matrix organization, angiogenesis and also to the regulation of cell migration. Moreover, we found that these four expression profiles contain genes known as markers of fetal Leydig cell precursors such as *Arx, Pdgfra, Gli2, Tcf21*, and *Ptch1* (**Figure 4B**) (Barsoum and Yao, 2011; Bhandari et al., 2012; Brennan et al., 2003; Cui et al., 2004; Inoue et al., 2015; Liu et al., 2016; Miyabayashi et al., 2013).

The transcriptome dynamics of the interstitial progenitor cell lineage reflect a change in cell identity, from multipotent epithelial cells expressing stem cell markers at E10.5 to steroidogenic precursor cells at the origin of fetal Leydig cells.

### Sertoli cell differentiation program is regulated by waves of transcription factors

Expression dynamics of key genes such as *Sry*, *Sox9*, *Fgf9* or *Dhh* is consistent with published reports and emphasize the complex gene expression kinetics at play during Sertoli cell lineage specification (**Figure 5A**). By applying the same procedure as we did for the progenitor cells, we identified the gene expression profiles driving Sertoli cell differentiation in the fetal testis from the E10.5 progenitor state prior to sex determination until E16.5. We identified 2319 genes differentially expressed as function of the pseudotime from E10.5 to E16.5 (*q-value*<0.05) and classified them into 13 gene profiles (**Figure 5B, Supplementary Data 6**). 1217 genes (profiles P7 to P13) characterize the Sertoli cells differentiation from E11.5, including *Sry*, *Sox9*, *Amh* and *Dhh*. We found transcription factors expressed in each of the expression profiles, with some presenting reproductive phenotypes in Mouse Genome Informatics (marked with an asterisk in **Figure 5B**) (MGI: http://www.informatics.jax.org) (Blake et al., 2017).

**Figure 5:**
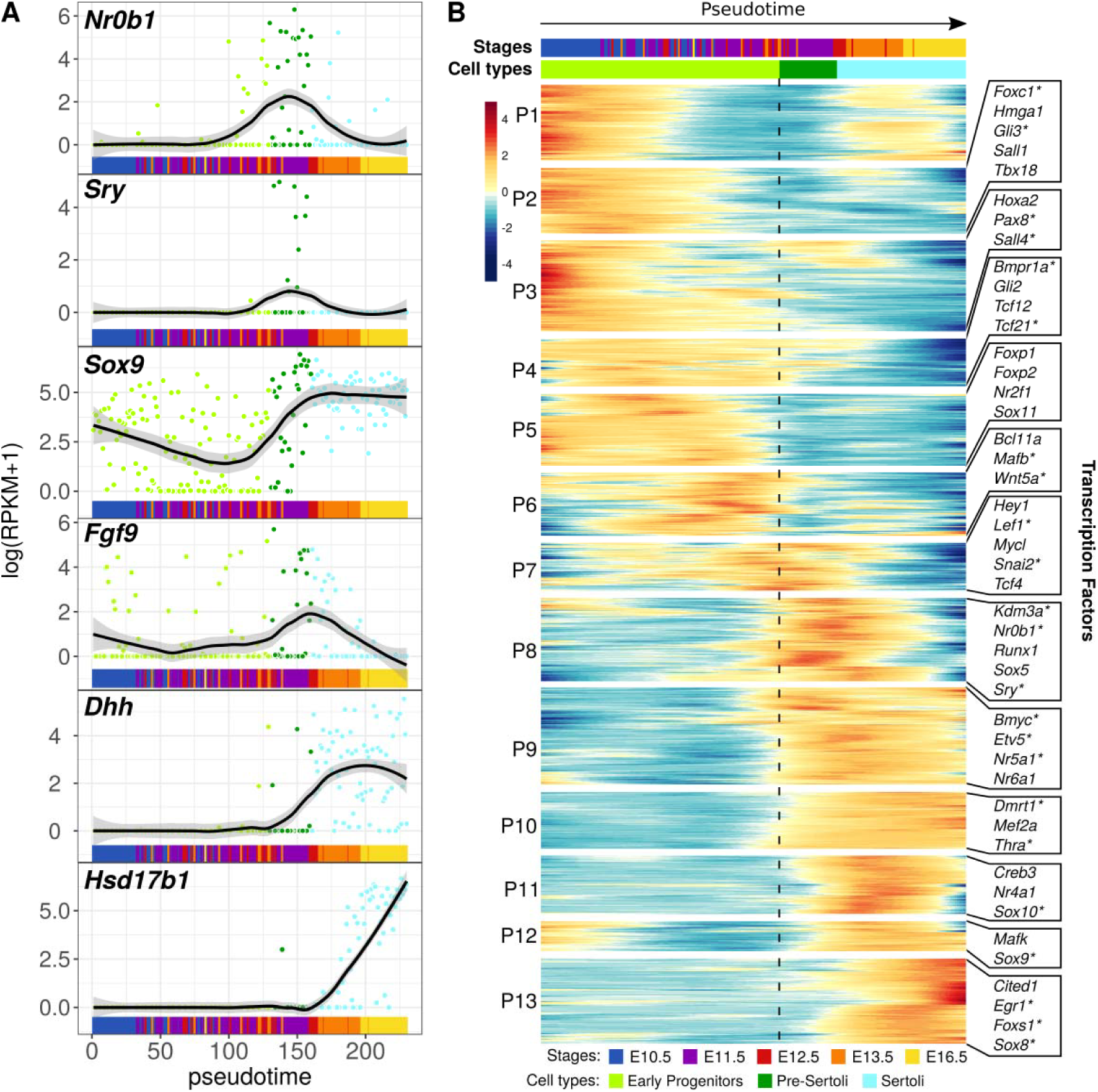
Figure 5: Sertoli cell transcriptome dynamics. (A) Expression profiles of known marker genes important for Sertoli cell differentiation. Each dot represent a cell, the solid line represents the loess regression, and the fade band is the 95% confidence interval of the model. (B) Heatmap of the normalized smoothed expression of genes differentially expressed as a function of the pseudotime. The vertical dash line represents the initiation of the Sertoli cell differentiation. On the right are listed some of the transcription factors expressed in each expression profile. Asterisks marks the ones presenting a reproductive phenotype in MGI.

Among the interesting expression profiles, we found 221 genes following the *Sry* narrow expression pattern at the onset of Sertoli cell differentiation (P8). They include *Kdm3a*, a H3K9 histone demethylase which directly regulate the expression of *Sry* by modifying the chromatin conformation upstream of *Sry* (Kuroki et al., 2013). *Nr0b1*, also known as *Dax1*, encodes a member of the orphan nuclear hormone receptor family. *Nr0b1* is considered as a pro-testis (Meeks et al., 2003) and anti-testis gene (Lovell-Badge et al., 1998).

In contrast, expression profiles P9, P10 and P13 display a long range activation in differentiating Sertoli cells. Among them, we find *Nr5a1* which co-regulates with *Sox9* the expression of *Amh* (expression profile 8, **Supplementary Data 6**) (Lasala et al., 2011), *Dmrt1* which has a key role in maintaining Sertoli cell fate and repressing female promoting signals (Lavery et al., 2012; Matson et al., 2011; Minkina et al., 2014), and *Gata4* which is required for testis differentiation and controls the expression of *Dmrt1* (Manuylov et al., 2011).

Our findings revealed that Sertoli cell differentiation is driven by a highly dynamic transcriptional program composed of several intermediate stages defined by peaks of co-expressed genes as well as activation and repression of genes. The transcriptomic signature of Sertoli cells evolves with time as a reflection of the differentiation status and evolving functions of these supporting cells during early testis development.

## Discussion

The classical model of gonadal sex determination states that the bipotential gonad contains at least two pre-established somatic cell lineages, the supporting and the steroidogenic cell lineages whose progenitors will ultimately differentiate into Sertoli and fetal Leydig cells, respectively (Svingen and Koopman, 2013). This model has been challenged toward a common progenitor, with the recent findings of Leydig-to-Sertoli trans-differentiation capacity (Zhang et al., 2015), and that both lineages derive from WT1^+^ cells present at E10.5 in the genital ridge (Liu et al 2016). However, the characterization of this multipotent somatic progenitor has so far remained elusive and the possibility that different populations of WT1^+^ progenitors carrying a supporting or steroidogenic fate, cannot be excluded.

Here we aimed at evaluating the heterogeneity of Nr5a1^+^ cells by establishing the transcriptomic identity of each individual cell present in the gonadal somatic compartment. Strikingly, we identify a single homogeneous multipotent NR5A1^+^ progenitor cell population in the bipotential gonad, whose transcriptomes progressively evolve and diverge toward the supporting and steroidogenic lineages from E11.5 onward. We cannot exclude the existence of additional rare NR5A1^+^ cell types, however our findings support the common progenitor identity hypothesis and demonstrate the absence of a cell lineage specific fate before sex determination, at least at the transcriptional level. The presence of a unique progenitor for both supporting and steroidogenic cell lineages raise the question of the mechanisms behind cell fate decision. We have showed that the progenitor cell lineage operates progressive transcriptomic changes through time, switching from a multipotent identity to a steroidogenic precursor identity. Before sex determination, cells express multipotent markers as well as *Wt1* and *Cbx2*, two co-factors necessary for *Sry* expression and Sertoli cell differentiation. From E11.5, when the Sertoli cells have differentiated, the progenitor cells lose expression of multiple genes including these two factors and as a consequence, might lose the capacity to differentiate as Sertoli cells.

The decision between a Sertoli fate and a progenitor fate during male sex determination remains elusive. However, we noticed the progenitor cells express genes involved in negative regulation of cell differentiation during the same time frame of Sertoli cell differentiation. Notably, we found *Sox11* which has been recently identified as a potential co-repressor of the pro-Sertoli gene *Sox9* (Zhao et al., 2017), as well as *Sfrp1*, a negative regulator of *Wnt* signaling pathway (Gauger et al., 2013). Possibly, the progenitor cells have a repression program allowing cells to escape the Sertoli cell fate and conserve a progenitor identity, similarly to a stem cell renewal process.

From E12.5, progenitor cells become restricted to the interstitial compartment of the testis and progressively lose *Wt1* gene expression, which is suspected to be the key regulator of the Sertoli cell differentiation and maintenance (Zhang et al., 2015). Conversely, they gradually express markers of fetal Leydig cell precursors, such as Arx, *Pdgfra*, and *Tcf21* (Brennan et al., 2003; Cui et al., 2004; Miyabayashi et al., 2013), and restrict their fate toward interstitial steroidogenic precursors at the origin of Leydig cells. The maintenance of the steroidogenic cell precursors is known to be regulated by a differentiation repression program via *Notch1* and *Tcf21.* This mechanism maintains a pool of fetal Leydig cell precursor cell population (Barsoum and Yao, 2010b; Tang et al., 2008).

We also noticed the interstitial progenitors express a significant amount of genes related to regulation of cell migration as well as angiogenesis. Our data suggest that the interstitial cells actively contribute to this cell migration and control the vascularization of the testis from E12.5, together with Sertoli cells. However, we found no clear evidence these cells are able to differentiate as endothelial cells.

Our results also reveal that Sertoli cell differentiation is mediated by a highly complex and dynamic transcriptional program composed of waves of transiently expressed genes. Our capacity to order the cells along a pseudotime reflecting their differentiation status allows the most accurate expression dynamics catalogue compared to published data based on pools of non-synchronous cells (Jameson et al., 2012; Nef et al., 2005). In particular, the narrow transient expression profile of numerous genes suggests they are required during a precise cellular differentiation stage but not for maintenance of cell identity. We found that both *Nr0b1* and *Sry* are expressed in E11.5 Pre-Sertoli cells during a very narrow window of time, confirming the hypothesis that both factors are critical for Sertoli cell differentiation and are required for *Sox9* upregulation (Eggers et al., 2014; Ludbrook and Harley, 2004). In contrast, a broad expression profile with a permanent activation in differentiating Sertoli cells might reveal a role in maintaining their identity. For example, the upregulation of *Sox9* and *Dmrt1* is consistent with their key role in maintaining Sertoli cell fate and repressing female promoting signals (Lavery et al., 2012; Matson et al., 2011; Minkina et al., 2014).

Testis formation is a complex developmental process requiring coordinated differentiation of multiple cell lineages. In the future, we expect single cell expression studies to transform our understanding of gonadal development and sex determination. In particular, a similar single-cell ovarian development analysis would provide a high resolution understanding of the mutually antagonist testis/ovarian genetic program at play during sex determination. Single-cell genomic will provide unparalleled insights into stem cell origin of the different cell lineages, distinguish the different stages of differentiation, characterize cell fate decisions and the regulatory mechanisms that govern the production of different cell types, and elucidate how different cell types work together to form a testis. We also expect that advances in single cell genomics such as Chip-seq, ATAC-seq or again Hi-C (Pott and Lieb, 2015; Ramani et al., 2017; Rotem et al., 2015) will facilitate the identification of epigenetic modifications and regions of open chromatin at the single cell level. In combination, such single cell approaches will serve as a basis for both understanding testis determination and its pathologies.

## Experimental Procedures

### Mouse strains and isolation of purified Nr5a1-GFP positive cells

Animals were housed and cared for according to the ethical guidelines of the Direction Générale de la Santé of the Canton de Genève (experimentation GE-122-15). The experiment has been performed using heterozygous Tg(Nr5a1-GFP) transgenic male mice (Stallings, 2002). We performed the experiments in independent duplicates for each embryonic stages, except for E10.5 where XX embryos were used as controls but not included in the study. At each relevant gestation days, Tg(Nr5a1-GFP) gonads were collected and dissociated. Several animals from different litters were pooled together to obtain enough material for the experiment (**Table S1**). GFP+ cells were sorted by fluorescent-active cell sorting (**Figure S1, Supplementary experimental procedures**).

### Tissue processing and immunological analyses

Embryos at relevant stages of development were collected, fixed in 4% paraformaldehyde overnight at 4°C, serially dehydrated and embedded in paraffin. Fluorescent images were acquired using a confocal laser scanning microscope and processed with ZEN software (**Supplementary experimental procedures**).

### Single-cell capture and cDNA libraries and sequencing

Cells were captured and processed using the C1 Autoprep System (96 well, small size IFC chip), following the official C1 protocol. Sequencing libraries were prepared using the Illumina Nextera XT DNA Sample Preparation kit using the modified protocol described in the C1 documentation. Cells were sequenced at an average of 15 million (**Supplementary experimental procedures**).

### Bioinformatics analysis

The computations were performed at the Vital-IT (http://www.vital-it.ch) Centre for high-performance computing of the SIB Swiss Institute of Bioinformatics. Data were analyzed with R version 3.4.0. Cell clustering was performed on the highly variable genes with HCPC, differential expression analysis with Monocle2, diffusion maps were computed with Destiny, and lineage reconstruction was performed with Slingshot (for details, see **Supplementary experimental procedures**).

## Author Contributions

I.S. and S.N. designed all experiments. I.S., Y.N., C.B. performed the experiments. I.S., and S.N. analyzed the data. J.E., S.E.A., E.T.D. and L.B.S. contributed to manuscript preparation. I.S., Y.N. and S.N. wrote the manuscript.

## Acknowledgments

Supported by grants from the Swiss SystemsX Interdisciplinary PhD Grant 51PHI0–141994 (to I.S.) and by the Département de l'Instruction Publique of the State of Geneva (to S.N.). We thank Luciana Romano and Deborah Penet for the sequencing, and Pascale Garlonne Ribaux and Françoise Kuhne for technical assistance. We thank also the members of the Nef and Dermitzakis laboratories for helpful discussion and critical reading of the manuscript, Cécile Gameiro from the flow cytometry facility for the cell sorting, and Didier Chollet, Brice Petit and Mylène Docquier from the iGE3 Genomics Platform for the single-cell capture.

